# Bird Song Learning is Mutually Beneficial for Tutee and Tutor

**DOI:** 10.1101/774216

**Authors:** Michael D. Beecher, Çağlar Akçay, S. Elizabeth Campbell

## Abstract

Song learning is generally assumed to be beneficial for a young songbird, but merely incidental, without costs or benefits, for the older song ‘tutors’. In the present study we contrast two mutually exclusive hypotheses about the tutor/tutee relationship: (1) that it is cooperative, or at least mutually tolerant, with tutor and tutee mutually benefiting from their relationship, *vs.* (2) that it is competitive, with tutor and tutee competing over territory, so that one or the other suffers negative fitness consequences of their relationship. In a field study of three consecutive cohorts of song sparrows (*Melospiza melodia*) we determined the older bird (primary tutor) from whom the young bird (tutee) learned most of his songs, and how long tutee and primary tutor survived subsequently. We found that the more songs a tutee learns from his primary tutor, the longer their mutual survival on their respective territories. This study provides the first evidence of a mutual benefit of bird song learning and teaching in nature.

## 1. Introduction

Young songbirds (oscine passerines) learn their songs from adult conspecifics, making songbirds one of a handful of taxa (including humans) that socially learn their vocalizations (Nowicki and Searcy, 2014). There are many notable similarities between bird song learning and human language learning (Doupe and Kuhl, 1999; Marler, 1970), but there is one striking difference: whereas in humans the tutors are typically parents, other relatives and other parties who gain by tutoring, in most songbirds song is learned from unrelated individuals who are most often territorial neighbours, and thus potential competitors for space, food, and mating opportunities (e.g., Beecher et al., 1994; Liu and Kroodsma, 2006; Payne, 1982; Wheelwright et al., 2008). Consequently, while human language learning is a cooperative, mutually beneficial process, one might expect the opposite to be true of bird song learning, at least in the common case where tutor and tutee are potential competitors. In this paper we consider the question of whether bird song learning is a mutually beneficial process or a competitive process in the song sparrow (*Melospiza melodia*).

It is generally assumed that young songbirds benefit from the particular songs they have learned. For starters, learning good conspecific song is prerequisite for acquiring a mate (Nowicki et al., 2002). Further, acquiring local songs of the neighbourhood to which they have dispersed has been shown to help young birds acquire mates and establish and keep a territory (Beecher et al., 2000b; O’Loghlen and Rothstein, 1995; Payne, 1982; Payne et al., 1988; Poesel et al., 2012; Wilson et al., 2000). These results suggest that sharing songs with your neighbours is advantageous and raises the question of whether the young bird’s tutor-neighbours might benefit as well.

To the best of our knowledge however, there is no evidence that older birds benefit by tutoring young birds. Indeed, it is often assumed that the older bird is simply singing in order to post its territory, interact with territorial neighbours or attract a mate, and that the young bird learns these songs as it eavesdrops on this singing (Beecher et al., 2007; Templeton et al., 2010). In this view the older bird’s song ‘tutoring’ is assumed to be incidental, not motivated. This scenario resembles the classical method for studying song learning, in which the young bird is ‘tutored’ by a loudspeaker playing recorded song in an isolation chamber (Marler and Peters, 1977; Marler and Peters, 1981). In a somewhat different scenario, song learning may involve direct singing interactions between the young bird and the older bird in an explicit competition over territory and mates. Learning the tutor’s songs could benefit the young bird in this competition, e.g., by allowing him to song match his rival (Akçay et al., 2017; Akçay et al., 2013; Burt et al., 2001). In both scenarios, the competition hypothesis (Beecher and Akçay, 2014) views tutor and tutee as rivals in a zero-sum game, in which case the survival of tutor and tutee should be negatively correlated, and the extent of the tutee’s learning of the tutor’s songs should reflect the intensity of this competition.

According to the mutual benefit hypothesis (Beecher and Akçay, 2014), in contrast, the tutor-tutee relationship is more cooperative than competitive. We can call the relationship an alliance, though it could simply be one of mutual tolerance. For the older bird an alliance gives him a young neighbour who is less threatening both in terms of territorial integrity and paternity confidence (in most songbirds, including song sparrows, extra-pair paternity is common and the extra-pair fathers are typically immediate neighbours (Akçay and Roughgarden, 2007; Hill et al., 2011; Hsu et al., 2015)). For the young bird the alliance gives him the ability to establish a territory more easily than he could otherwise. Although being allies does not necessarily require song sharing between the tutee and tutor, song sharing may facilitate this relationship, e.g., by signalling to others that these neighbours are allies. Evidence for a functional role of shared vocalizations for coordination and affiliation comes from group-living animals in which shared vocalizations positively influence group cohesion and bonds (King and McGregor, 2016; Tyack, 2008).

In our sedentary population of song sparrows in the northwestern United States the processes of song learning and territory establishment are correlated. A male song sparrow disperses from his natal area at about a month of age, and acquires a repertoire of about 9 songs (e.g., Figure S1) during his first year of life, learning from unrelated adults in the area where he attempts to establish his territory (Akçay et al., 2014; Beecher et al., 1994; Nordby et al., 1999). The bird crystallizes his song repertoire by the time he is fully territorial sometime between January and March of the next spring. He will keep his territory and his song repertoire (unchanged) for the rest of his life (average 3 years).

We tested the mutual benefit and competition hypotheses by measuring the survival of a tutee and his primary song tutor as a function of how many of his songs the tutee had learned from the tutor. Song sparrows in this population typically have a primary tutor from whom they learn anywhere from 30% to 100% of their songs (in the present data set the mean percentage was 55%, Figure S2). We reason that if song learning is mutually beneficial for the song tutor and the young song learner, then the more of the tutor-neighbour’s songs the young bird learns, the more years the two will coexist on their territories (their mutual survival). This hypothesis assumes that the full benefit of song sharing between tutor and tutee occurs only so long as both remain on territory as neighbours. In contrast, according to the competition hypothesis, the more of his songs the young bird learns from the older tutor, the fewer years the two will coexist on territory (their mutual survival). This hypothesis assumes that how much the young bird learns from this tutor reflects the degree of their territorial competition and that the presence of one of them makes the future survival of the other one less likely (Akçay et al., 2017).

## 2. Methods

### (a) Study Site and Subjects

Our study site is a 200-ha undeveloped city park bordering Puget Sound in Seattle, Washington, with ca. 100–150 colour-banded males on territories per year. Males hold territories year-round in this population (Nordby et al., 1999). The song-learning period starts early in the bird’s natal summer, about a month after the bird hatches (usually in April, May or June), and persists into autumn and the next spring (Nordby et al., 2001; Nulty et al., 2010). We captured song sparrows with mist nets either passively or using playback and banded them with U.S. Fish and Wildlife Service metal bands and a unique combination of three coloured leg bands that allowed visual identification of individuals later.

Our cohorts were 34 birds that hatched in 2009, 24 that hatched in 2010 and 20 that hatched in 2011. We recorded the song repertoires of these 78 young birds and compared them to the recorded repertoires of approximately 120 older birds in each year who were their potential tutors. We considered all adult males present in June of a young bird’s hatch year as potential tutors.

### (b) Recording, Song Analysis and Censusing

Male song sparrows sing 7 to 11 song types (full range 6-13), and deliver these songs in bouts, singing one song type a number of times before switching to another song type (‘eventual variety’). We recorded the full repertoire of all the tutees and potential tutors (see supplementary methods for details on sampling and equipment). Song learning analyses were carried out as described in (Akçay et al., 2014; Nordby et al., 1999, and supplementary materials). Briefly, for each male we printed out spectrograms of several variations of each song type (Syrinx, John Burt, www.syrinxpc.com). Each tutee song was compared to all of the songs of the potential tutors by three or four judges independently, after which the judges arrived at a consensus sheet.

To determine survival on territory, we surveyed all territories in the park from 2009 onwards at least once per month. We assumed a male has lost his territory if we could not attract him with playback and if another bird was now observed defending the male’s territory. We also checked the neighbouring territories to ensure the male had not shifted his territory. Song sparrows in our population rarely move more than one territory away from where they first established their territory, and they rarely regain a territory after losing it to another male (Arcese, 1989; Beecher, 2008). We characterized survival in terms of the number of breeding seasons survived (survival to at least April).

### (c) Data analyses

We ran mixed-effect models with cohort as a random factor, proportion of repertoire learned from the primary tutor as the fixed factor and the following variables as response variables in separate models: tutor survival, tutee survival and mutual survival of tutor and tutee, the latter defined as the time period when both birds were alive. Because we were using the same predictor variable in this set of models, we corrected for multiple comparisons by taking alpha to be 0.05/3= 0.016. We also present the coefficients of determination (R^2^) from each model (Nakagawa et al., 2017).

In our previous studies, we found that tutor survival to the late stages of song learning (i.e., into the first spring) has a direct, proximate effect on song learning of the tutee, in that tutees generally learn more from tutors who survive into their first spring (Akçay et al., 2014; Nordby et al., 1999). This direct, proximate effect of tutor survival on tutee song learning should not be confounded with the indirect, ultimate survival benefit of song sharing between tutor and tutee. Since our question was did the amount learned from the primary tutor predict survival, we excluded those cases where we knew survival of the primary tutor (into the first spring) predicted the amount learned from him, i.e., where cause and effect were potentially reversed. We therefore repeated the above analyses by excluding all primary tutors who did not survive into the tutee’s first spring (n=11). Finally, we also asked whether the repertoire size of the tutees predicted their survival on the territory as might be expected if larger repertoires are an indicator of higher quality individuals (Pfaff et al., 2007) using mixed-models as above but repertoire size as the fixed factor.

## 3. Results

The models (Table 1) showed that proportion of repertoire learned from the primary tutor positively predicted the primary tutor’s survival, the tutee’s survival, and their mutual survival in the full dataset. After excluding those tutors that did not survive into the first spring of the tutee, only the tutee’s survival and mutual survival showed a positive relationship with proportion learned from primary tutor. The effect was stronger for mutual survival and only in that model did the p-value remain significant after correction for multiple comparisons. These results are consistent with the mutual benefit hypothesis and inconsistent with the competition hypothesis. Repertoire size of the tutees did not predict survival (supplementary materials).

**Table 1.** Results of the linear mixed models for each of the three survival variables (tutor survival, tutee survival and mutual survival) with proportion learned from primary tutor as the predictor variable: coefficients (and standard errors), t-values and the associated p values, and the marginal R^2^s (variance explained by the fixed factor). All models have cohort as a random factor and proportion learned from the primary tutor as the only fixed factor. The right side of the table shows the subset of data in which primary tutors that did not survive into the first spring of the tutee are excluded.

|  | **All data (n=78)** |  |  |  | **Primary tutors dying before Jan 1 excluded (n=67)** |  |  |  |
| --- | --- | --- | --- | --- | --- | --- | --- | --- |
| **Survival** | Coefficient (SE) | t | p | R<sup>2</sup> (marginal) | Coefficient (SE) | t | p | R<sup>2</sup> (marginal) |
| **tutor** | **2.44 (0.95)** | **2.56** | **0.013** | **0.076** | 1.45 (1.01) | 1.44 | 0.16 | 0.029 |
| **tutee** | 1.80 (0.92) | 1.95 | 0.055 | 0.044 | 2.19 (1.01) | 2.17 | 0.034 | 0.062 |
| **mutual** | **2.20 (0.58)** | **3.81** | **0.0003** | **0.13** | **1.67 (0.61)** | **2.72** | **0.008** | **0.078** |

## 4. Discussion

Prior to this study, we knew that a song sparrow in his first breeding season benefits by sharing songs with his neighbours (some of whom will have been his tutors); indeed the degree of this song sharing is the best predictor of the young bird’s long-term retention of his territory (Beecher et al., 2000b). In the present study we find that the proportion of a tutee’s song repertoire that he learned from his primary tutor – and which they share for so long as they both survive as territorial neighbours – predicts both the survival of the tutee and the mutual survival of the tutor and tutee on their territories. The effect on mutual survival is stronger than that on the survival of the tutee or tutor alone, and suggests that the benefits of song sharing accrue only so long as both individuals are alive and present on territory.

Parallel results have been found in indigo buntings, *Passerina cyanea*, a territorial songbird in which males sing a single, complex song (Payne, 1982; Payne et al., 1988). The investigators focused on indigo buntings in their first breeding season, and found that young males who shared their song with an older neighbour (the presumptive song tutor) had greater breeding success than did young males who shared with a distant male (the presumptive song tutor) or with no one at all in the population. That is, breeding success was dependent on the coexistence of both the young bird and his presumptive tutor on adjacent territories, another parallel with the present study. Finally, trends for adults were similar (adults sharing with a neighbour fared better), but tutor-tutee relationships were generally unknown in these cases, thus these trends can only suggest the possibility of a mutual benefit for tutor and tutee.

### (a) An Alliance Between Tutor and Tutee?

The mutual survival benefit we have observed may arise from an alliance developed by the tutee and his primary tutor where shared songs function as a ‘badge’ indicating familiarity and a cooperative relationship (Abolins-Abols et al., 2016; Brown and Farabaugh, 1997; Brown et al., 1988; Farabaugh et al., 1988; Wilson and Vehrencamp, 2001). The theoretical rationale for why territorial neighbours would benefit from forming a defensive coalition has been laid out by Getty (Getty, 1987). Briefly, it is that once two neighbours have agreed on the location of their border, it would be more costly for either of them to have to negotiate a new boundary with a new neighbour than to mutually defend their existing boundary against an intruder, and it would be more costly for the intruder to fight this coalition than to try and find an undefended or poorly-defended territory elsewhere. Support for this view has been found in the territorial red-winged blackbird (*Agelaius phoeniceus*). In this species the presence of familiar neighbours has been shown to increase the fitness of a territory holder (Beletsky and Orians, 1989). There is evidence that this advantage may result in part from neighbours forming coalitions for nest defence (Olendorf et al., 2004a). Moreover, when presented with neighbour or stranger song at their territory centre, males respond more aggressively to neighbours, also consistent with Getty’s argument that the neighbour’s defection from the alliance warrants an especially strong retaliation (Getty, 1987; Olendorf et al., 2004b).

We suggest a simple extension of Getty’s argument: when a song sparrow with an established territory loses an immediate neighbour, it could be advantageous for him to groom a juvenile male for that territory. Social interaction has been shown repeatedly to be a stronger stimulus for song learning than simple overhearing of song (review in Beecher, 2017), so the older bird could teach the young bird many of his songs by counter-singing with him while behaving less aggressively toward him than he would toward an adult prospecting for a territory. In an earlier study on likely tutor-tutee neighbours (i.e., one bird in his first breeding season, the other bird older, and the two sharing half or more of their repertoires), we found that they typically do type-match one another (reply with the same song type) early in the breeding season, but repertoire-match one another (reply with a different shared song type) later in the breeding season (long-term neighbours typically repertoire-match at any point in the year) (Beecher et al., 2000a). In theory, type-matching is the optimal way for the tutor to shape the young bird’s developing songs to more closely match his.

### (b) Are Primary Tutors “Dear Enemies”?

Although on occasion we have seen song sparrow neighbours jointly defend against intruders, we do not have strong evidence that a young bird and a strong primary tutor typically form a cooperative alliance. It is possible that the relationship is only one of mutual tolerance. This mutual tolerance, usually called a Dear Enemy relationship, has been found in many territorial animals, including song sparrows (Fisher, 1954; Getty, 1987; Jaeger, 1981; Temeles, 1994; Ydenberg et al., 1988). The Dear Enemy relationship may vary in the degree of tolerance, and while it may fall short of a true alliance, it is still clearly preferable to a fully competitive relationship.

Might primary tutors, particularly those that had strong influence on the tutee’s song repertoire, be particularly dear enemies, i.e., more mutually tolerant than most neighbouring birds? Two earlier studies on our study population have given conflicting results. In one playback study we simulated an intrusion onto the territory of the young bird before the onset of his first breeding season (in February or early April). We used self-song as a proxy for tutor song because (1) the song learning analysis can only be carried out after the breeding season, and (2) since the young bird would have only recently learned the song from one or more of his tutor-neighbours, he would likely perceive it as from one of them. We found that the young birds responded less aggressively to the intrusion if they had a strong primary tutor (Ç. Akçay & M. D. Beecher, unpublished data). Although this was a territorial intrusion, it was still prior to the breeding season, at a time when a young bird’s tolerance of his primary tutor might permit this. Adult song sparrows do tolerate young birds on their territory in the natal summer and fall, and these associations do sometimes continue into early spring (Templeton et al., 2012a; Templeton et al., 2012b).

On the negative side, in another playback experiment we simulated an intrusion by a former tutor (no longer alive) onto the territory of his former tutee (now in his second or later breeding season). In this case we found that birds responded more aggressively to stronger former tutors (Akçay et al., 2017). On the face of it, the results of the study described above are consistent with the mutual benefit hypothesis, while the results of this study are consistent with the competition hypothesis. However, Getty’s theoretical argument above suggests an alternative interpretation of this second study, namely that the stronger response to former tutors, especially the most influential ones, represents an especially strong retaliation against a neighbour who has broken the Dear Enemy agreement by the territorial intrusion. Prior playback experiments in our population have shown that a single simulated intrusion by a neighbour is sufficient to break down the Dear Enemy effect not only for the victim of the intrusion (Akçay et al., 2009), but even for other neighbours who simply observed the intrusion (Akçay et al., 2010). These results suggest that song sparrows play a tit-for-tat like strategy to maintain their Dear Enemy relationships but that an intrusion by a former ally violating the agreement is taken as a greater threat – a ‘double cross’ – and so evokes a stronger response, than an intrusion by either a stranger or even a familiar bird with whom the subject has a weaker dear enemy relationship.

This perspective predicts that in the normal course of territoriality, the Dear Enemy effect should be stronger between a bird and his primary tutor than between the bird and his other neighbours who were not tutors, or were weaker tutors. Indeed, there is often variation in the degree of Dear Enemy effect: some neighbours are dearer enemies than others (Stoddard, 1996). For instance, in another resident western population of song sparrows, Wilson and Vehrencamp (Wilson and Vehrencamp, 2001) found that the more songs a bird shared with his neighbour, the less aggressive his response to playback of the neighbour’s song at their mutual territory boundary (cf. Payne, 1983). Although the authors did not identify tutor-tutee relationships in that study, the degree of sharing suggests that at least some of these pairs were likely tutee and primary tutor (about half the subjects shared at least half their songs with the tested neighbour, which in our population usually indicates a primary tutor).

The final and perhaps most convincing piece of evidence for the mutual benefit hypothesis, is our observation that young song sparrows rarely replace their primary tutors (Akçay et al., 2014; Nordby et al., 1999) as should be relatively common if learning songs from a particular older bird is part of a strategy to take over his territory. Instead tutees are typically adjacent neighbours of their primary tutors, while the birds the tutees typically replace are either non-tutors or weak tutors (responsible for 1 or 2 songs at most). Thus while we have minimal evidence that the relationship of tutee and his primary tutor is explicitly cooperative (e.g., that they routinely co-defend against intruders or predators), the totality of the evidence suggests that it is least mutually tolerant, which would be consistent with our finding that it is mutually beneficial.

### (c) Bird song learning as an instance of teaching in non-human animals

The present finding of a long-term mutual benefit of song learning for tutor-tutee pairs suggests that bird song learning might be viewed as an instance of teaching in animals (Caro and Hauser, 1992; Hoppitt et al., 2008; Thornton and Raihani, 2010). The commonly accepted criteria for demonstrating teaching in non-human animals are that (1) teachers should modify their behaviour in the presence of the learner, (2) this change in behaviour should result in no immediate benefit to the teacher, and (3) the learner should acquire a behaviour quicker or better as a result (Caro and Hauser, 1992). Bird song tutoring would fit all three criteria if in fact the older bird reduces his usual aggression when a young bird appears on his territory (e.g. (Templeton et al., 2012a)), increases his counter-singing with the young bird, and the young bird learns many of the older bird’s songs. Direct observational evidence on these points will be needed to evaluate whether this instance of bird song tutoring qualifies as teaching.

## 5. Conclusion

In summary, we found that mutual survival of tutors and tutees is positively associated with the degree to which the latter learn their songs from the former. Our results suggest that learning many songs from a particular tutor-neighbour is beneficial for both parties, suggesting a cooperative or at least mutually tolerant relationship. Direct evidence for such a relationship will require further studies focusing on the behaviours that potentially underlie the mutual benefit effect. These would include the behaviours we have described that could be considered teaching by the older birds, and subsequent cooperative behaviours, such as joint defence of mutual boundaries.

## Acknowledgements

We thank Discovery Park for hosting our research, and Christopher Templeton, Veronica Reed and Saethra Darling for their help with field work and song learning analyses. This research was supported by grants from NSF (IOS-0733991) and the University of Washington Royalty Research Fund.

## Figures and Tables

**Figure 1.**
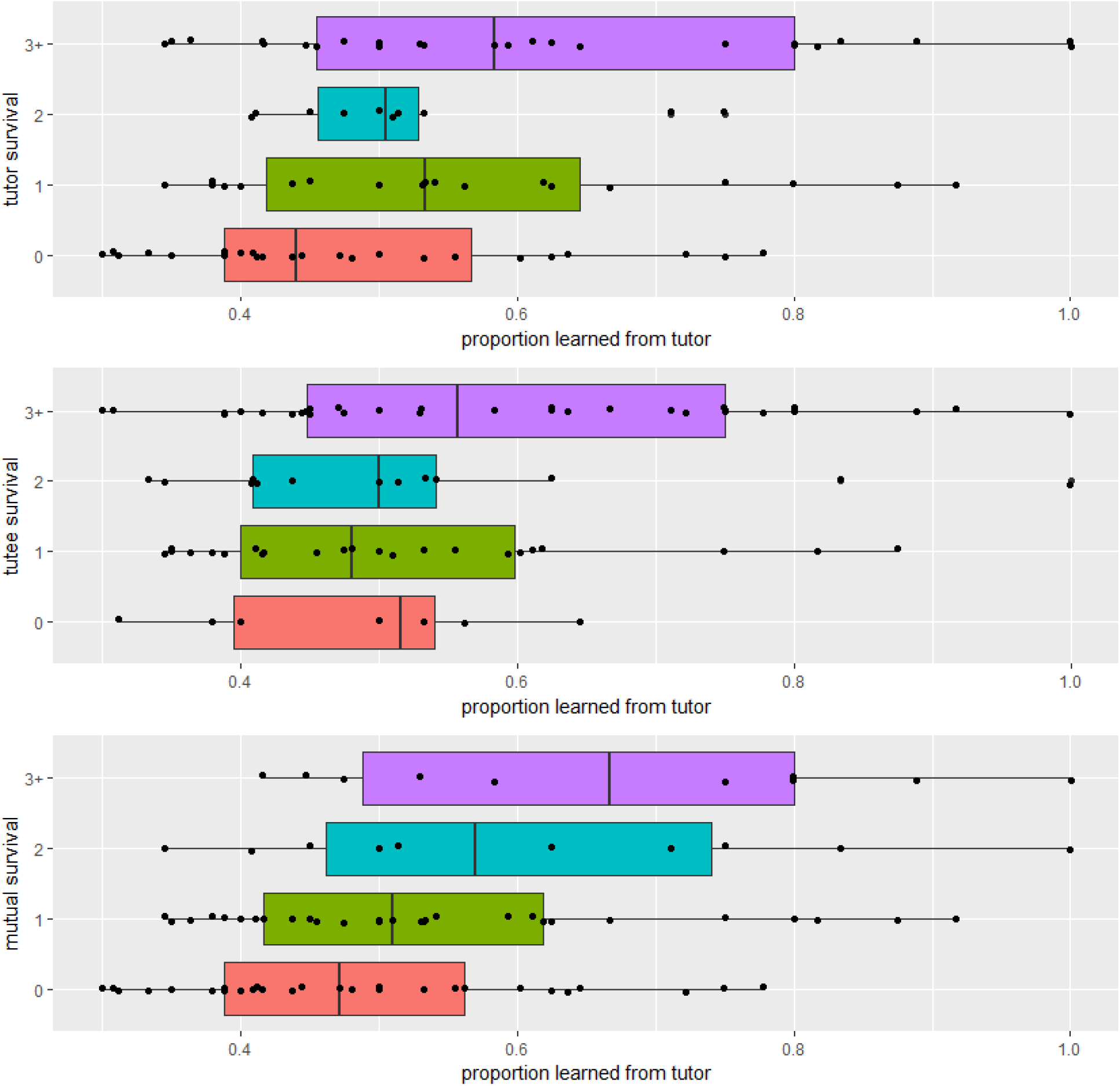
Years (breeding season) primary tutor, tutee or both survived a function of the proportion of the tutee’s song repertoire that was learned from the primary tutor.

## Literature cited

Abolins-Abols M, Hope SF, Ketterson ED, 2016. Effect of acute stressor on reproductive behavior differs between urban and rural birds. Ecology and evolution 6:6546-6555.

Akçay Ç, Campbell SE, Beecher MD, 2017. Good tutors are not dear enemies in song sparrows. Anim Behav 129:223-228. doi: https://doi.org/10.1016/j.anbehav.2017.05.026.

Akçay Ç, Campbell SE, Reed VA, Beecher MD, 2014. Song sparrows do not learn more songs from aggressive tutors. Anim Behav 94:151-159.

Akçay Ç, Reed VA, Campbell SE, Templeton CN, Beecher MD, 2010. Indirect reciprocity: song sparrows distrust aggressive neighbors based on eavesdropping. Anim Behav 80:1041-1047.

Akçay Ç, Tom ME, Campbell SE, Beecher MD, 2013. Song type matching is an honest early threat signal in a hierarchical animal communication system. Proc R Soc Lond, Ser B: Biol Sci 280:20122517. doi: http://dx.doi.org/10.1098/rspb.2012.2517.

Akçay Ç, Wood WE, Searcy WA, Templeton CN, Campbell SE, Beecher MD, 2009. Good neighbour, bad neighbour: song sparrows retaliate against aggressive rivals. Anim Behav 78:97-102.

Akçay E, Roughgarden J, 2007. Extra-pair paternity in birds: review of genetic benefits. Evol Ecol Res 9:855-868.

Arcese P, 1989. Territory acquisition and loss in male song sparrows. Anim Behav 37:45-55.

Beecher MD, 2008. Function and mechanisms of song learning in song sparrows. Advances in the study of behavior 38:167-225.

Beecher MD, 2017. Birdsong learning as a social process. Anim Behav 124:233-246. doi: https://doi.org/10.1016/j.anbehav.2016.09.001.

Beecher MD, Akçay Ç, 2014. Friends and enemies: how social dynamics shape communication and song learning in song sparrows. In: Yakusawa K, editor. Animal Behavior Santa Barbara, CA: Praeger.

Beecher MD, Burt JM, O’Loghlen AL, Templeton CN, Campbell SE, 2007. Bird song learning in an eavesdropping context. Animal Behavior 73:929-935.

Beecher MD, Campbell SE, Burt JM, Hill CE, Nordby JC, 2000a. Song type matching between neighboring song sparrows. Animal Behavior 59:21-27.

Beecher MD, Campbell SE, Nordby JC, 2000b. Territory tenure in song sparrows is related to song sharing with neighbors, but not to repertoire size Animal Behavior 59:29-37.

Beecher MD, Campbell SE, Stoddard PK, 1994. Correlation of song learning and territory establishment strategies in the song sparrow. Proc Natl Acad Sci USA 91:1450-1454.

Beletsky LD, Orians GH, 1989. Familiar Neighbors Enhance Breeding Success in Birds. Proceedings of the National Academy of Science 86 7933-7936.

Brown ED, Farabaugh SM, 1997. What birds with complex social relationships can tell us about vocal learning: vocal sharing in avian groups. In: Snowdon CT, Hausberger M, editors. Social Influences on Vocal Development Cambridge: Cambridge University Press. p. 98-127.

Brown ED, Farabaugh SM, Veltman CJ, 1988. Song sharing in a group-living songbird, the Australian magpie, *Gymnorhina tibicen*: Part I. Vocal sharing within and among social groups. Behaviour 104:1-28.

Burt JM, Campbell SE, Beecher MD, 2001. Song type matching as threat: a test using interactive playback. Anim Behav 62.

Caro TM, Hauser MD, 1992. Is there teaching in nonhuman animals? The Quarterly Review of Biology 67:151-174.

Doupe AJ, Kuhl PK, 1999. Birdsong and human speech: Common themes and mechanisms. Annu Rev Neurosci 22:567-631.

Farabaugh SM, Brown ED, Veltman CJ, 1988. Song sharing in a group-living songbird, the Australian magpie: Part II. Vocal sharing between territorial neighbors, within and between geographic regions, and between sexes. Behaviour 104:105-125.

Fisher JB, 1954. Evolution and bird sociality. In: Huxley J, Hardy AC, Ford EB, editors. Evolution as process London: Allen & Unwin. p. 71-83.

Getty T, 1987. Dear enemies and the prisoner’s dilemma: why should territorial neighbors form defensive coalitions? Am Zool 27:327-336.

Hill CE, Akçay Ç, Campbell SE, Beecher MD, 2011. Extrapair paternity, song and genetic quality in song sparrows. Behav Ecol 22:73-81.

Hoppitt WJ, Brown GR, Kendal R, Rendell L, Thornton A, Webster MM, Laland KN, 2008. Lessons from animal teaching. Trends Ecol Evol 23:486-493.

Hsu YH, Schroeder J, Winney I, Burke T, Nakagawa S, 2015. Are extra-pair males different from cuckolded males? A case study and a meta-analytic examination. Mol Ecol 24:1558-1571.

Jaeger RG, 1981. Dear enemy recognition and the costs of aggression between salamanders. Am Nat 117:962-979.

King SL, McGregor PK, 2016. Vocal matching: the what, the why and the how. Biol Lett 12:20160666.

Liu W-C, Kroodsma DE, 2006. Song learning by chipping sparrows: When, where, and from whom. Condor 108:509-517.

Marler P, 1970. Birdsong and speech development: could there be parallels? Am Sci 58:669-673.

Marler P, Peters S, 1977. Selective vocal learning in a sparrow. Science 198:519-521.

Marler P, Peters S, 1981. Sparrows learn adult song and more from memory. Science 213:780-782.

Nakagawa S, Johnson PC, Schielzeth H, 2017. The coefficient of determination R 2 and intra-class correlation coefficient from generalized linear mixed-effects models revisited and expanded. Journal of the Royal Society Interface 14:20170213.

Nordby JC, Campbell SE, Beecher MD, 1999. Ecological correlates of song learning in song sparrows. Behav Ecol 10:287-297.

Nordby JC, Campbell SE, Beecher MD, 2001. Late song learning in song sparrows. Anim Behav 61:835-846.

Nowicki S, Searcy W-A, Peters S, 2002. Quality of song learning affects female response to male bird song. Proceedings of the Royal Society Biological Sciences Series B.

Nowicki S, Searcy WA, 2014. The evolution of vocal learning. Curr Opin Neurobiol 28:48-53. doi: https://doi.org/10.1016/j.conb.2014.06.007.

Nulty B, Burt JM, Akçay Ç, Templeton CN, Elizabeth Campbell S, Beecher MD, 2010. Song learning in song sparrows: relative importance of autumn vs. spring tutoring. Ethology 116:653-661.

O’Loghlen AL, Rothstein SI, 1995. Culturally correct song dialects are correlated with male age and female song preferences in wild populations of brown-headed cowbirds. Behavioral Ecology & Sociobiology 36:251-259.

Olendorf R, Getty T, Scribner K, 2004a. Cooperative nest defence in red–winged blackbirds: reciprocal altruism, kinship or by–product mutualism? Proceedings of the Royal Society of London Series B: Biological Sciences 271:177-182.

Olendorf R, Getty T, Scribner K, Robinson SK, 2004b. Male red-winged blackbirds distrust unreliable and sexually attractive neighbors. Proc R Soc Lond, Ser B: Biol Sci 271:1033-1038.

Payne RB, 1982. Ecological consequences of song matching: breeding success and intraspecific song mimicry in indigo buntings. Ecology 63:401-411.

Payne RB, 1983. The social context of song mimicry: song-matching dialects in indigo buntings *(Passerina cyanea)*. Anim Behav 31:788-805.

Payne RB, Payne LL, Doehlert SM, 1988. Biological and cultural success of song memes in indigo buntings. Ecology 69:104-117.

Pfaff JA, Zanette L, MacDougall-Shackleton SA, MacDougall-Shackleton EA, 2007. Song repertoire size varies with HVC volume and is indicative of male quality in song sparrows (Melospiza melodia). Proceedings of the Royal Society B-Biological Sciences 274:2035-2040.

Poesel A, Nelson DA, Gibbs HL, 2012. Song sharing correlates with social but not extrapair mating success in the white-crowned sparrow. Behav Ecol 23:627-634.

Stoddard PK, 1996. Vocal recognition of neighbors by territorial passerines.. In: Kroodsma DE, Miller DE, editors. Ecology and evolution of acoustic communication in birds Ithaca, New York: Cornell University Press. p. 356-374.

Temeles EJ, 1994. The role of neighbors in territorial systems: when are they ’dear enemies’? Animal Behavior 47:339-350.

Templeton CN, Akçay Ç, Campbell SE, Beecher MD, 2010. Juvenile sparrows preferentially eavesdrop on adult song interactions. Proceedings of the Royal Society B: Biological Sciences 277:447-453. doi: 10.1098/rspb.2009.1491.

Templeton CN, Campbell SE, Beecher MD, 2012a. Territorial song sparrows tolerate juveniles during the early song-learning phase. Behav Ecol 23:916-923.

Templeton CN, Reed VA, Campbell SE, Beecher MD, 2012b. Spatial movements and social networks in juvenile male song sparrows. Behav Ecol 23:141-152.

Thornton A, Raihani NJ, 2010. Identifying teaching in wild animals. Learn Behav 38:297-309. doi: 10.3758/lb.38.3.297.

Tyack PL, 2008. Convergence of calls as animals form social bonds, active compensation for noisy communication channels, and the evolution of vocal learning in mammals. Journal of Comparative Psychology 122:319.

Wheelwright NT, Swett MB, Levin II, Kroodsma DE, Freeman-Gallant CR, Williams H, 2008. The influence of different tutor types on song learning in a natural bird population. Anim Behav 75:1479-1493.

Wilson PL, Towner MC, Vehrencamp SL, 2000. Survival and song-type sharing in a sedentary subspecies of the song sparrow. The Condor 102:355-363.

Wilson PL, Vehrencamp SL, 2001. A test of the deceptive mimicry hypothesis in song-sharing song sparrows. Anim Behav 62:1197-1205.

Ydenberg RC, Giraldeau L-A, Falls JB, 1988. Neighbours, strangers and the asymmetric war of attrition. Anim Behav 36:343-347.

